# Reusable and Sensitive Detection Method for Exonuclease III Activity by DNB Nanoarrays Based on cPAS Sequencing Technology

**DOI:** 10.1101/2020.03.24.002956

**Authors:** Ying Chen, Hui Wang, Jin Yang, Huanming Yang, Radoje Drmanac, Chongjun Xu, Wenwei Zhang

## Abstract

In this article, we have designed a sensitive and recycled DNB (DNA nanoball) nanoarrays sequencing complex structures based on BGISEQ-500RS sequencer for the monitoring performance of Exo III activity. In the shortage of Exo III, the effective number ratio of DNB would be captured by an optical system due to one fluorescent. In contrast, in the presence of Exo III, some DNB would disappear or discard from the fields of the optical system by fluorescence extinction and uncleaned fluorescent, respectively. As a result, the effective number of DNB of this strategy was relative to the concentration of Exo III. For Exo III, our strategy showed a highly sensitive linear response in the low detection range of 0.01 U/mL to 0.5 U/mL, with detection limits below 0.01 U/mL. With the comparison between DNB nanoarrays and other fluorescent sensors, this study possessed superior sensitivity, selectivity, and reusability, accompanying with the low cost and simple setup.

## 1. Introduction

Exonucleases are DNA enzymes that participate in a hydrolyzing reaction which break phosphodiester bonds at the end of a polynucleotide chain, and cleave one nucleotide at a time. The exonuclease family is concerned with the sustenance of cellular metabolism and physiological processes, such as aiding DNA proofreading and sustaining the stability of the genome [1]. Exonuclease III (Exo III) belongs to the exonuclease family with 3’-termini to 5’ -termini exonuclease activities, that works by stepwise removal of mononucleotide from the 3’-OH termini of doublestranded DNA when the substrates are recessed or blunt, and the 3’-terminus does not require any specific recognition site and presents lower activities on single-stranded DNA or duplex DNAs with a protruding 3’-terminus, acts an indispensable role in some cellular and physiological processes, and is also imperative to preserving the stability of the genome[2]. In the process of DNA replication, Exo III owns the requisite capacities in DNA proofreading, stimulating gene recombination reactions, and repairing DNA double-strand breaks [3, 4], undertaking the precision of the DNA replication process and preserving the mutation rates in a cell [5, 6]. Studies [7] have described that the deficiencies in both over and low expression of 3’ to 5’ end exonucleases can begin cells to deliver inaccurate transcription and misleading translation, which subsequently increases the risk to cancerization, especially under prolonged stress. Therefore, investigations for the detection of Exo III are significant because they can be used as a detector for diseases. Unquestionably, it is completely urgent to develop well-performed analytical methods that can produce a rapid and precise measurement of the activity of 3’→5’exonucleases in a complex biological milieu [8].

Ordinarily, conventional approaches for 3’ end to 5’ end exonuclease activity detection are based on gel electrophoresis and radioactively labeled DNA probes [9]. But, these traditional methods are characterized by time-consuming, tedious steps, inherent security concerns, and more higher costs [10, 11]. To overcome the restrictions mentioned above, scholars have attempted numerous studies on biosensing systems for 3’→5’exonuclease activity. For instance, a label-free detection method based on K^+^-induced G-quadruplex and ThT dye for Exo III by using a DNA hairpin probe has advantages as simple and low cost [12]. The device of graphene oxide-based fluorescence assay promotes the detection sensitivity of Exo III from 0.01U to 0.05U [13]. The Tb 3+-induced G-quadruplex conjugates form a rapid and label-free fluorescent “turn-on” assay for Exo III detection [14]. The interaction between SYBR Green I and double-strand DNA has stimulated a novel assay for realtime monitoring of the 3’→5’exonucleases [15]. In Jiang’ group, they used the principle of long-range resonance energy transfer between graphene oxide and ethidium bromide, developed a simple label-free fluorometric scheme for DNA exonuclease activity [16]. The multiple exonuclease detection method is created by a triple color fluorescent probe which can serve as a lab-on-a-DNA-molecule for multiple exonucleases or restriction endonucleases detection synchronous [17]. Nevertheless, the above-mentioned techniques displayed one or more disadvantages, such as insensitive, time-consuming or uneconomical.

DNB (DNA nanoball) nanoarrays provides the array of fluorescent signals in DNB sequencing technology, which can be captured by a sequencer optical system. Besides, through the effective number ratio of DNB analysis, uncleaned fluorescent DNB will be discarded at last. This study intends to develop a new economy and rapid Exo III detection method through DNB sequencing technology. In this work, the DNB sequencing complex (DSC), as the substrate of Exo III, is the 3’-recessed end doublestrand DNA structure that can polymerize the blocked-fluorescently-labeled dNTPs and exhibit signals. This strategy can detect Exo III activity with detection limits below 0.01 U/mL. With the comparison between DNB nanoarrays and other fluorescent sensors, this assay occupies superior sensitivity, selectivity, and reusability, accompanying with the low cost and simple setup.

## 2. Materials and Preparations

### 2.1 Materials

In this part, we introduce materials or reagents used in this project. MGIEasy PCR-Free DNA Library Prep Set (Cat. No. 1000013452) for ssDNA (circle DNA) library construction. BGISEQ-500RS Make & Load Reagent 4rxn kits (Cat.No.1 000008235) are to prepare DNB nanoarrays. BGISEQ-500RS PE100 Sequencing Reagent kit which consists of wash buffer, reaction buffer and dye marked dNTPs (Cat.no.1000008229) can provide fluorescent signals for detection. All of the abovementioned kits are purchased from MGI (BGI, China). Exo III with a specific activity of 100 U/ml, Exonuclease I (10 U/ml), T4 PNK (10 U/ml) and 10 × reaction buffer we used in this research are obtained from MGI (BGI, China) too. 0.5M EDTA (pH8.0) buy from Sigma. TE buffer is from Invitrogen (pH8.0, 10mM tris, and 1mM EDTA). Deoxynucleotide Solution (dATP, dCTP, dGTP, dTTP) and Lambda Exo buy from NEB. Ultrapure water (18.2MΩcm^−1^) is utilized during all the experiments.

### 2.2 Oligonucleotides

The Ecoli genomics (Bacterial species: ATCC-8739) obtained from the microbiology lab of the China national gene bank (CNGB). Oligonucleotides we used in the experiments are showed in Table 1, which are synthesized by Beijing Liuhe (BGI Techsolutions), and dissolved in TE buffer and stored at −20 °C for further use.

**Table 1.**
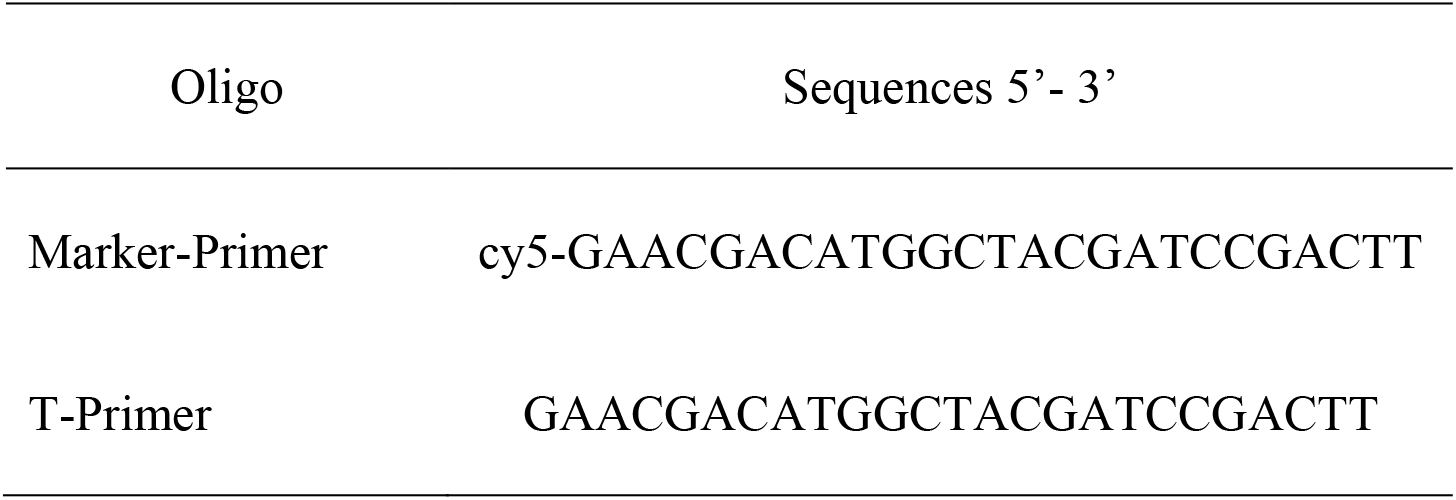
Oligonucleotides information

### 2.3 Preparations

We follow the instructions of MGIEasy PCR-Free DNA Library kit mentioned above, which are for ssDNA construction, and get 2.15ng/uL ssDNA with the main size on 320nt. The concentration of 20min rolling circle amplification (RCA) products is 8ng/ul when ssDNA inputs are 40fmol. The DNB is loaded on a nanoarray flow cell by following instructions of BGISEQ-500RS Make & Load kit.

## 3. Results and discussion

In this section, we have described the experimental procedures and results. A schematic illustration of the main experimental process of Exo III activity detection is shown in Figure 1, which is executed on BGISEQ-500RS sequencer with BGI-cPAS sequencing technology. In single-terminal sequencing of BGI-cPAS sequencing technology, the sequencing primers or reads sequences combined on the DNB template that could form a 3’-blunt end or 3’-recessed end of dsDNA and produce signals when the blocked and fluorescently-labeled dNTPs were polymerized. This DNB sequencing complex (DSC) are the substrate of Exo III, which only catalyze the stepwise removal of mononucleotides from 3’-termini of dsDNA. While the presence of Exo III, DSC structure can be digested and released free DNB, that cannot be detected by the sequencer optics system. When the DNB template rehybridize primers, each DNB would be read again through the specific fluorescence signal. Additionally, Exo III activity probably be judged from signal to effective number ratio of DNB (ESR) too. Therefore, this propositional design allows the sensitivity to the Exo III activity.

**Figure 1.**
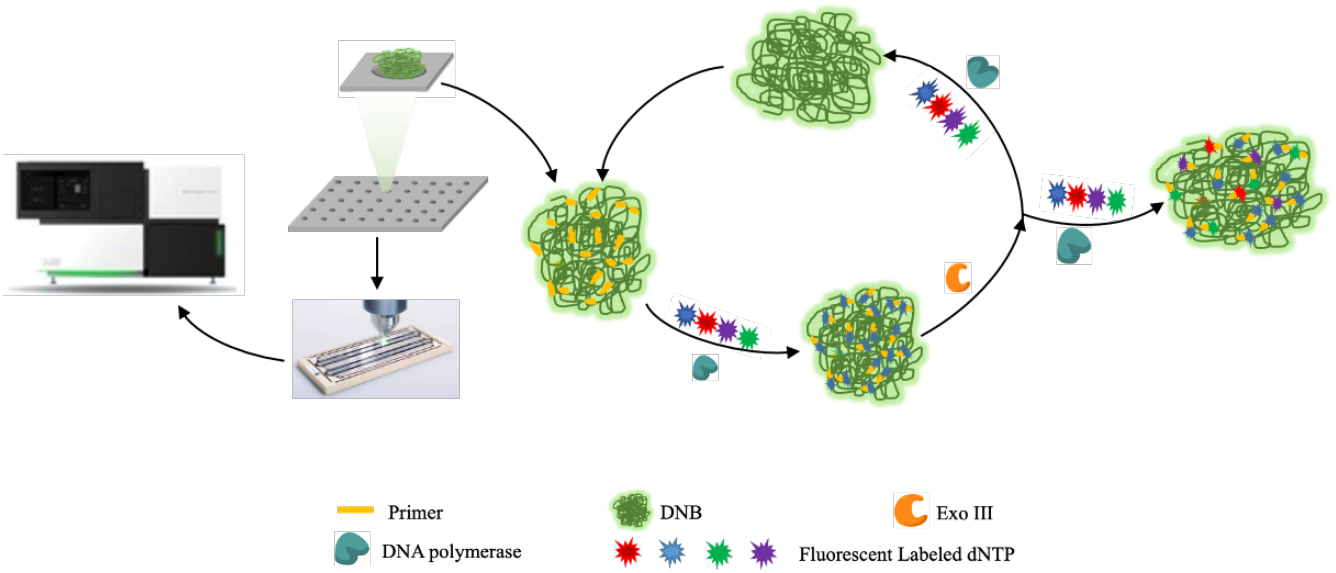
Schematic illustration of the reusable and sensitively Exo III detection with based on cPAS sequencing technology and BGISEQ-500RS sequencer system.

### 3.1 Feasibility of Exo III detection

To verify the feasibility of the current strategy, a simple scheme is performed. The 3’-recessed end of dsDNA we obtain from the hybridization of Cy5 labeled primer and DNB complex. For the better hydrolyzation to this substrate of Exo III, we incubate it on a heater 30min with 37 degrees. We take photos on BGISEQ-500RS sequencer before and after Exo III treatment. Results, as shown in Figure 2, are visible that the 3’-recessed end of dsDNA could be identified by Exo III, while DNB is not recognized. The Exo III hydrolyzes dye Cy5 labeled primers from DNB, which make it impossible to see any DNB in the field. The new Cy5 labeled primers lead DNB exposed to the optics detection system.

**Figure 2.**
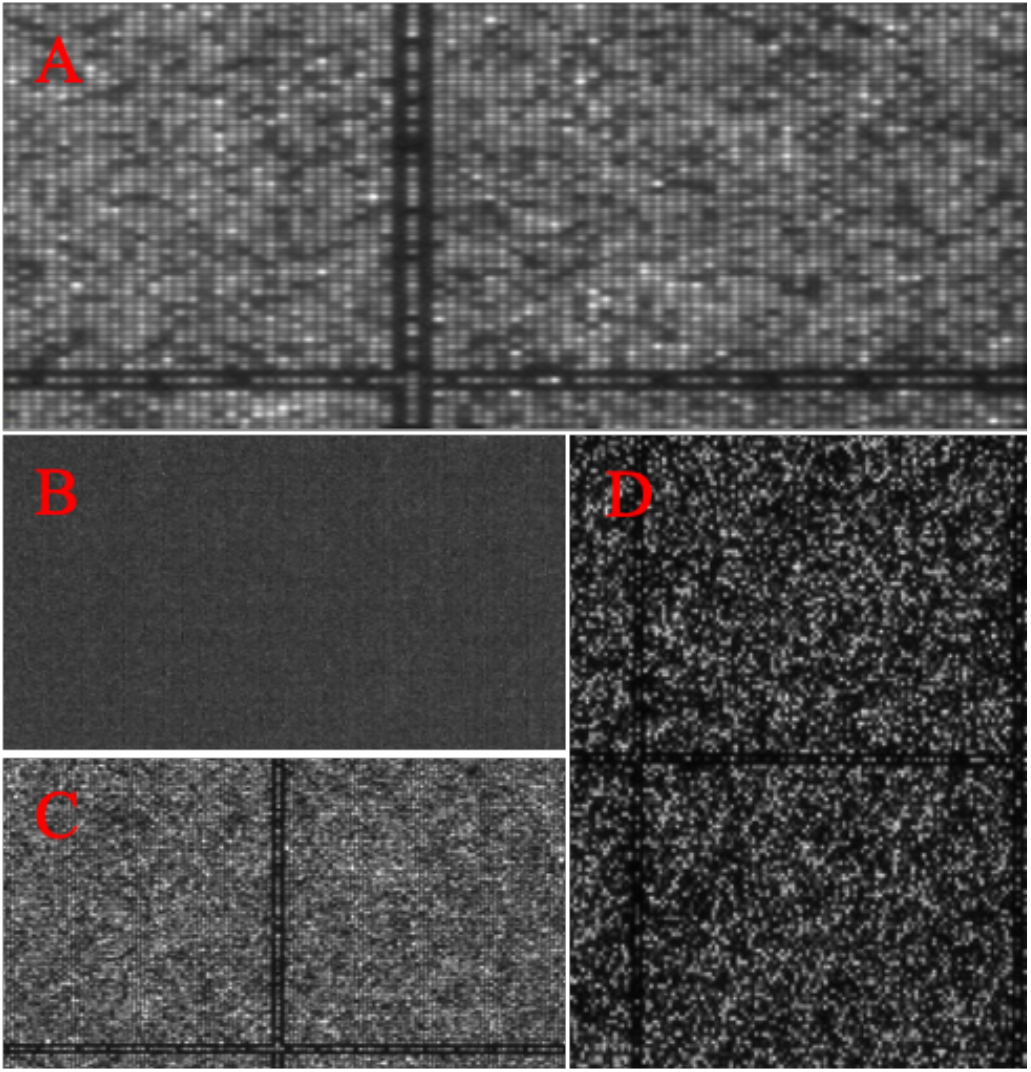
The DNB imaging results from BGISEQ-500RS sequencer. A) The DNB formed a 3’-recessed end structure with Cy5 dye labeled primer (Marker-Primer), made DNB visible through the BGISEQ-500RS sequencer; B) The DNB image results after EXO III treatment, fluorescence signals lost due to the 3’-recessed end structure in A were identified by Exo III; C) DNB image reproduce again through rehybridization of Marker-Primers and DNB; D) DNB image results of Channel A when T-Primer and fluorescently-labeled dNTPs were used

### 3.2 Specificity investigation

Many DNA enzymes have some effects in common with Exo III and play essential roles in multiple biological processes. For some examples, the Exo I is the enzyme to cut down single-strand DNA from 3’ end to 5’ end [18]. Lambda Exo removes mononucleotides of 5’ end from a duplex DNA[19]. T4 PNK is hunting for 5’ end phosphorylation of DNA/RNA for subsequent ligation[20, 21].To validate the specificity of methods for Exo III detection, we measure fluorescence signal of the nanoarray DNBs by the sequencer towards those DNA enzymes which treated at 37 degrees 10 min with a concentration of 2U/mL. As shown in Figure.3, except for Exo III, no other DNA enzymes we used in this research could generate significant changes in fluorescence intensity and DNB number-efficiency ratio, which is an indication of excellent specificity for Exo III detection by using this technique.

**Figure 3.**
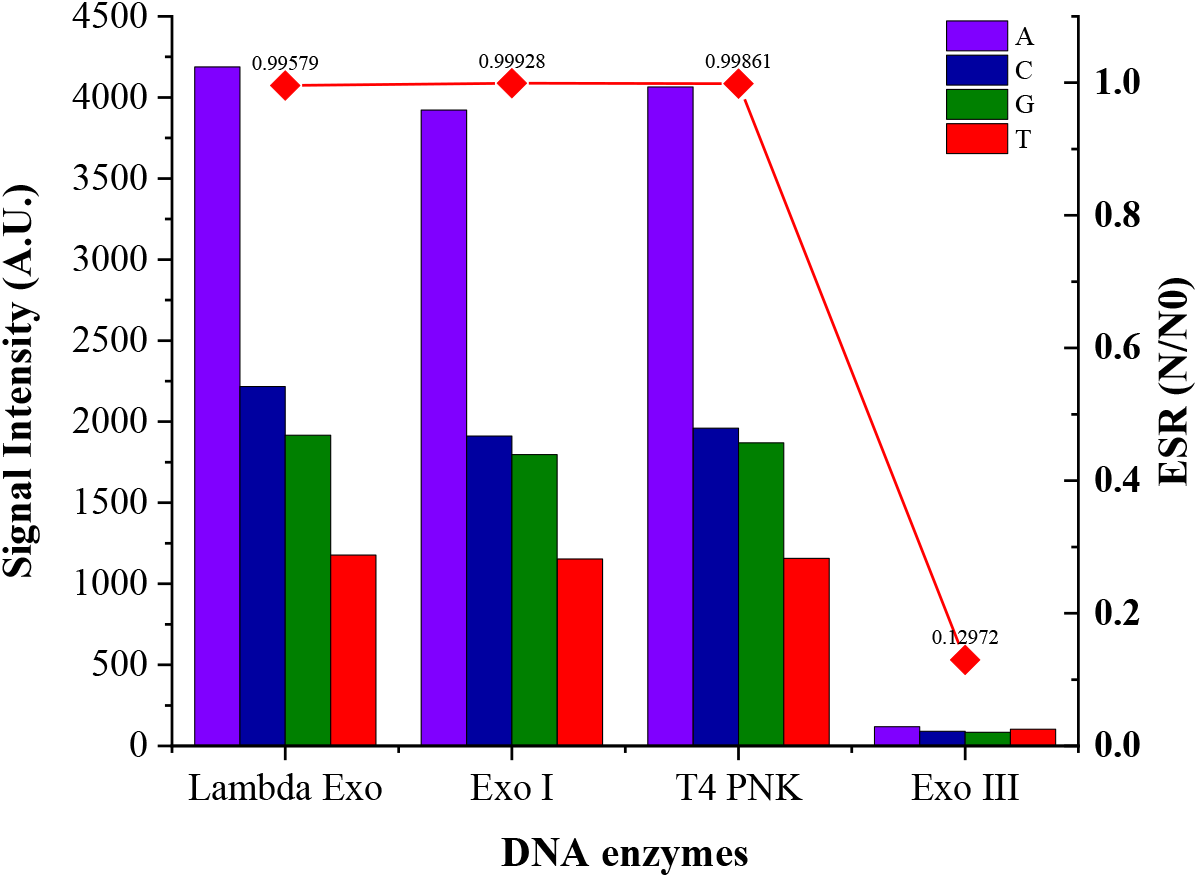
Specificity for Exo III detection. Fluorescence intensity of four dNTPs and DNB number efficiency ratio in different DNA enzyme.

### 3.3 Sensibility for Exo III

To survey the sensitivity of our proposal method towards Exo III, we methodically evaluate the relationships between ESR values and Exo III performance by varying concentrations of Exo III from 0 to 2 unit/mL (0, 0.01, 0.05, 0.1, 0.25, 0.5, 1, 1.5, 2 U/mL) for 37 degrees, following by incubation with 10min. We extract the fluorescence signal of four kinds of dNTPs from photos acquired by the BGISEQ-500RS sequencer optical system, then calculate the ESR in that detection area. The corresponding ESR values change with the Exo III concentration presented in Figure 4. Apparently, it increases gradually as the concentration of Exo III increase, with a highly sensitive linear response at the range from 0.01 U/mL to 0.5 U/mL (details were in the inset of Figure 4). The calibration curve of ESR at the linear range concentration shows the regression equation of Y=0.0091X+0.9579 with a correlation coefficient of 0.9612. Y axis is the ESR ratio values, and X axis presents the concentration of Exo III. Note, while the Exo III concentration is too high, the ESR ratio may decrease due to enzyme residues. Buffer washing also perhaps cause a slight dropping of the ESR ratio, which probably led by the quenching of our fluorescent dyes.

**Figure 4.**
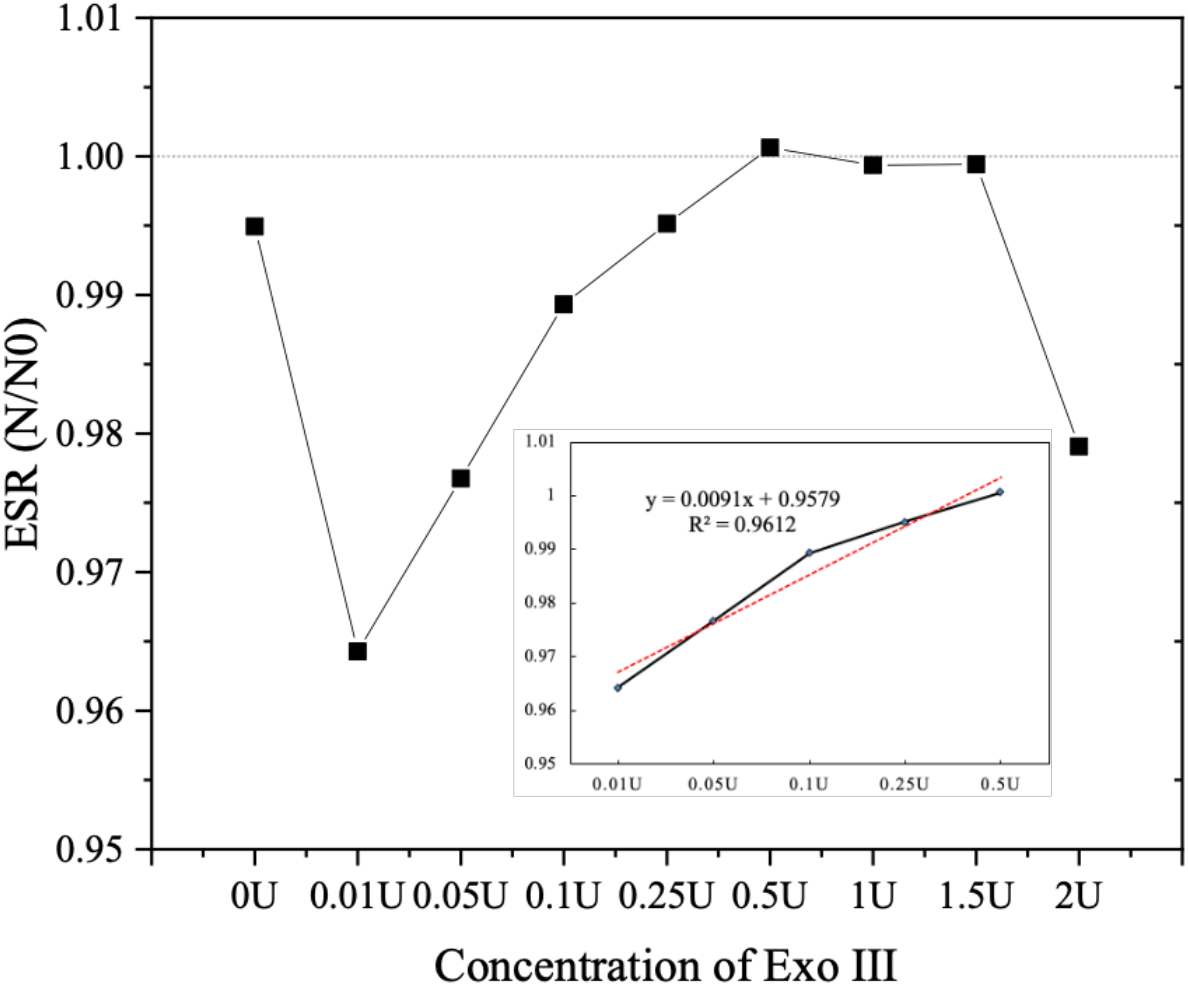
DNB number efficiency values of different concentration of Exo III. The inset showed a linear relationship between the DNB number efficiency values versus Exo III concentrations.

The detection limit of our proposed research plan is estimated to be lower than 0.01 U/mL, which is comparable to or lower than most of the Exo III detection methods reported previously (Table 2). Taken together, the results of this study demonstrate that DNB nanoarrays based on BGISEQ-500RS sequencer system might provide a simple approach for the accurate quantification of Exo III activity by possessing superior sensitivity, selectivity, and reusability, accompanying with the low cost and simple setup.

**Table 2.** Comparisons of the present Exo III detection method with other fluorescent methods

| Method | Detection range (U/mL) | Detection limit (U/mL) | Response time | Reference |
| --- | --- | --- | --- | --- |
| Graphene oxide | 5–120 | 1 | 60min | [16] |
| SYBR Green I | 1–200 | 0.7 | 110min | [15] |
| Tb3+ | 5–100 | 0.8 | 45min | [14] |
| CuNPs | 0.02–10 | 0.02 | 35min | [2] |
| ThT | 0–10 | 0.5 | 30min | [12] |
| GO/HP | 0.01-0.5 | 0.01U | 20min | [13] |
| DNB nanoarrays | 0.01-0.5 | < 0.01U | 10min | This work |

To further recognize the relationships between the excision capability of Exo III and the double helix length of DNA in this DNB nanoarray method, DNB doublestrand length of 30bp, 60bp, 90bp, 120bp, 150bp, 180bp, 210bp and 240bp are generated respectively by using a single-end sequencing (SE sequencing) on cPAS BGISEQ-500RS sequencer system. In this section, the working concentration and temperature of Exo III are 0.5U/mL and 37degree respectively, after 10 minutes, T-primers rehybridize on the DNB template. Then, the camera of the BGISEQ-500RS sequencer collects fluorescence intensities from each nanoarray DNBs when cPAS reactions are performing. As Figure 5 shows, the length range of double-helix DNA is below 60bp, the recover fluorescence intensities variations are not significant. While the DNA double-helix length is from 60bp to 120bp, fluorescence signals are weakened, but not obviously. As the length range is over 120bp, we can observe the recover signals keep decaying. These phenomena present that the double-strand length of a DNB sequencing complex (DSC) can affect the efficiency of Exo III which make some signals of a part of DNB copy numbers are invalid and regarded during signal analysis by the sequencer software.

**Figure 5.**
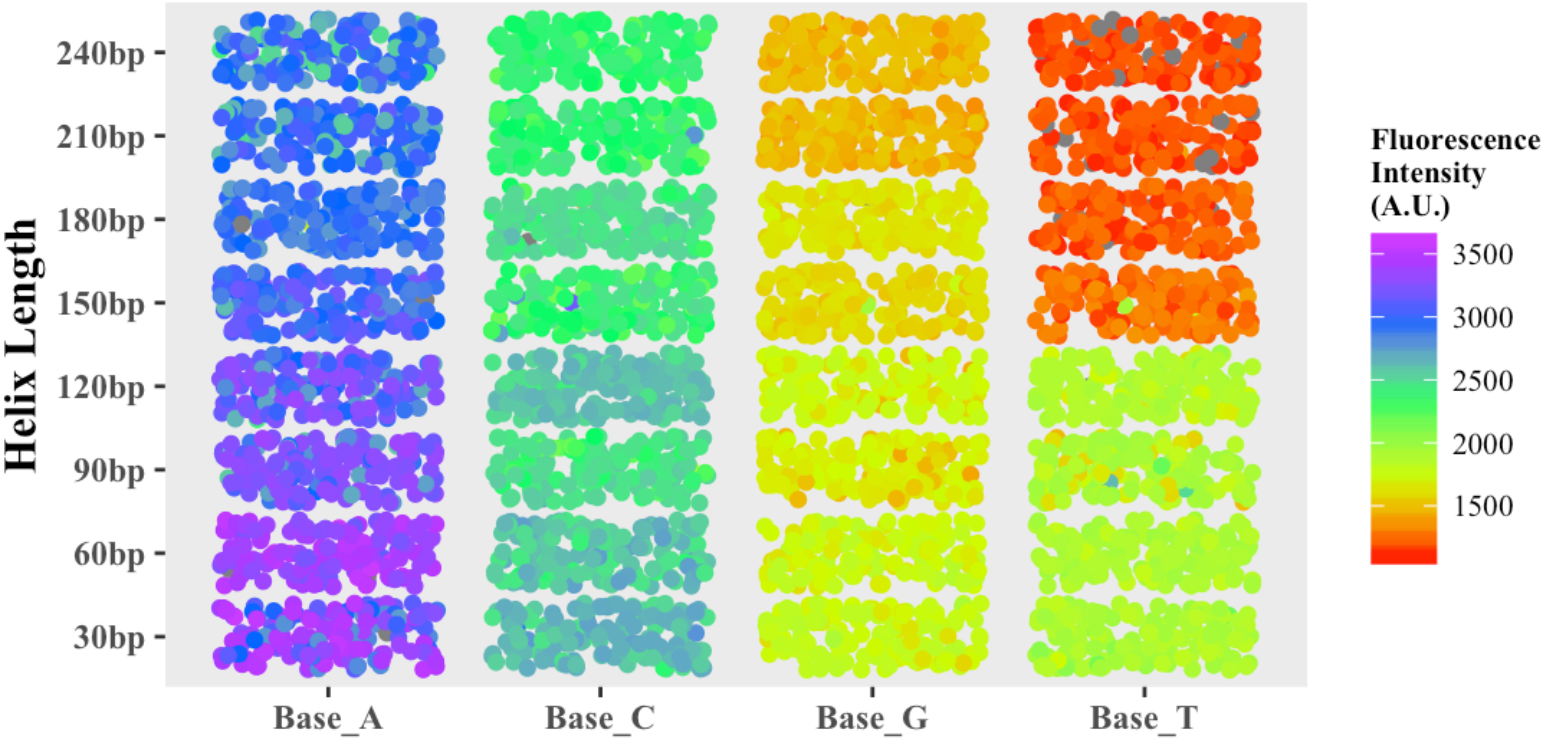
The recover fluorescence intensities of four fluorescently-labeled dNTPs after Exo III excision reactions at the different DNB double-strand helix length.

### 3.4 Reusability verification

For the repeatability of this technology for Exo III detection, we collect the performance of ESR to experiment frequency. Our experimental process in this part is that DNB is captured by sequencer first under fluorescently-labeled dNTPs. Then, Exo III is used to remove fluorescently-labeled sequences from DNB. Third, new primers and fluorescently-labeled dNTPs led DNB visible and form the new DNB sequencing complex structures which are the substrate of Exo III. All the experiments of this part conduct at the same optimum conditions, that is, Exo III is working 10min at 37 degrees with a concentration of 0.5U. 1X reaction buffer is employed in control experiments. As shown in Figure 6, the ESR values keep much stabilized even we did five times of experiments on one DNB nanoarrays flow cell, indicating that DNB sequencing technology with BGISEQ-500RS sequencer for Exo III detection is characterized by recycled, sensitively and specific. Y represents the ESR ratio, and X is the number of experiments by Exo III working.

**Figure 6.**
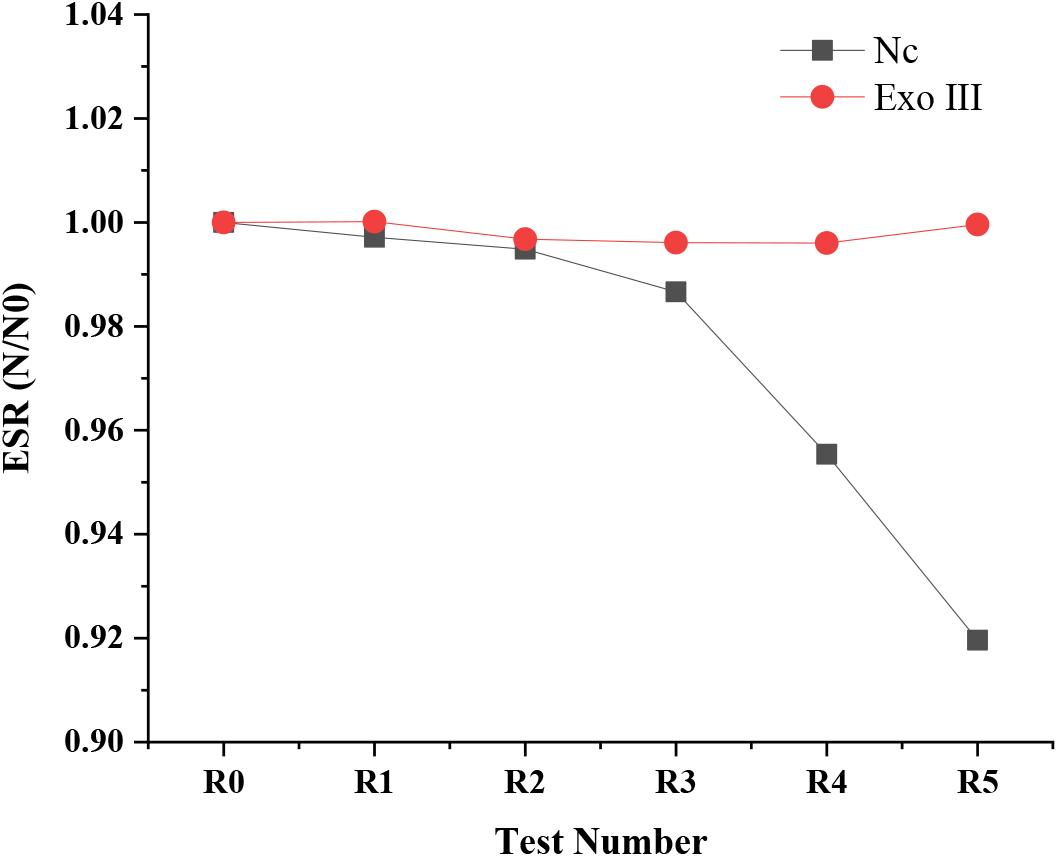
DNB number efficiency values of reusability tests. X axis was the number of experiments loops.

## 4. Conclusions

In summary, a highly sensitive and reusability project for the detection of Exo III activity has developed, which depends on the DNB sequencing technology based on digestion from the 3’-recessed end of DNB sequencing structures by Exo III. The method prefers good sensibility to Exo III with a detection limit of 0.01 U/mL. Furthermore, compared with other reported studies, this strategy has traits of repeat use, simplification experimental techniques, and none DNA design, which could be more feasible and lower cost. Concurrently, this method exhibits high sensitivity and selectivity for the detection of Exo III, which is anticipated to be employed for disease diagnosis and therapy.

## Declaration of competing interest

The authors have declared no conflict of interest.

## Acknowledgments

This work was supported by the Enterprise project of BGI-Shenzhen, China.

## References

[1] K.M. Burkin, O.L. Bodulev, A.V. Gribas, I.Y. Sakharov, One-step label-free chemiluminescent assay for determination of exonuclease III activity towards hairpin oligonucleotides, Enzyme Microb Technol, 131 (2019) 109419. https://doi.org/10.1016/j.enzmictec.2019.109419

[2] J. Ge, Z.-Z. Dong, D.-M. Bai, L. Zhang, Y.-L. Hu, D.-Y. Ji, Z.-H. Li, A novel label-free fluorescent molecular beacon for the detection of 3’–5’ exonuclease enzymatic activity using DNA-templated copper nanoclusters, New Journal of Chemistry, 41 (2017) 9718–9723. https://doi.org/10.1039/C7NJ01761H

[3] L. Song, M. Chaudhuri, C.W. Knopf, D.S. Parris, Contribution of the 3’-to 5’-exonuclease activity of herpes simplex virus type 1 DNA polymerase to the fidelity of DNA synthesis, Journal of Biological Chemistry, 279 (2004) 18535–18543. doi: 10.1074/jbc.M309848200

[4] C.D. Mol, C.-F. Kuo, M.M. Thayer, R.P. Cunningham, J.A. Tainer, Structure and function of the multifunctional DNA-repair enzyme exonuclease III, Nature, 374 (1995) 381–386. doi: 10.1038/374381a0

[5] A.M. Whitaker, T.S. Flynn, B.D. Freudenthal, Molecular snapshots of APE1 proofreading mismatches and removing DNA damage, Nature communications, 9 (2018) 1–11. https://doi.org/10.1038/s41467-017-02175-y

[6] Y.-C. Chen, C.-L. Li, Y.-Y. Hsiao, Y. Duh, H.S. Yuan, Structure and function of TatD exonuclease in DNA repair, Nucleic acids research, 42 (2014) 10776–10785. https://doi.org/10.1093/nar/gku732

[7] I.V. Shevelev, U. Hübscher, The 3’–5’ exonucleases, Nature Reviews Molecular Cell Biology, 3 (2002) 364–376. https://doi.org/10.1038/nrm804

[8] C.-J. Wang, W. Lam, S. Bussom, H.-M. Chang, Y.-C. Cheng, TREX1 acts in degrading damaged DNA from drug-treated tumor cells, DNA repair, 8 (2009) 1179–1189. https://doi.org/10.1016/j.dnarep.2009.06.006

[9] M. Brucet, J. Querol-Audí, K. Bertlik, J. Lloberas, I. Fita, A. Celada, Structural and biochemical studies of TREX1 inhibition by metals. Identification of a new active histidine conserved in DEDDh exonucleases, Protein Science, 17 (2008) 2059–2069. https://doi.org/10.1110/ps.036426.108

[10] A.V. Nimonkar, A.Z. Özsoy, J. Genschel, P. Modrich, S.C. Kowalczykowski, Human exonuclease 1 and BLM helicase interact to resect DNA and initiate DNA repair, Proceedings of the National Academy of Sciences, 105 (2008) 16906–16911. https://doi.org/10.1073/pnas.0809380105

[11] D.A. Lehtinen, S. Harvey, M.J. Mulcahy, T. Hollis, F.W. Perrino, The TREX1 double-stranded DNA degradation activity is defective in dominant mutations associated with autoimmune disease, Journal of biological chemistry, 283 (2008) 31649–31656. doi: 10.1074/jbc.M806155200

[12] X. Jiang, H. Liu, F.Y. Khusbu, C. Ma, A. Ping, Q. Zhang, K. Wu, M. Chen, Label-free detection of exonuclease III activity and its inhibition based on DNA hairpin probe, Anal Biochem, 555 (2018) 55–58. https://doi.org/10.1016/j.ab.2018.06.014

[13] X. Liu, Y. Wu, X. Wu, J.X. Zhao, A graphene oxide-based fluorescence assay for the sensitive detection of DNA exonuclease enzymatic activity, Analyst, 144 (2019) 6231–6239. https://doi.org/10.1039/C9AN01283D

[14] W. Yang, Y. Ruan, W. Wu, P. Chen, L. Xu, F. Fu, A “turn-on” and label-free fluorescent assay for the rapid detection of exonuclease III activity based on Tb 3+-induced G-quadruplex conjugates, Analytical and bioanalytical chemistry, 406 (2014) 4535–4540. https://doi.org/10.1007/s00216-014-7830-8

[15] M. Xu, B. Li, Label-free fluorescence strategy for sensitive detection of exonuclease activity using SYBR Green I as probe, Spectrochimica Acta Part A: Molecular and Biomolecular Spectroscopy, 151 (2015) 22–26. https://doi.org/10.1016/j.saa.2015.06.052

[16] Y. Jiang, J. Tian, S. Chen, Y. Zhao, Y. Wang, S. Zhao, A Graphene Oxide–Based Sensing Platform for The Label-free Assay of DNA Sequence and Exonuclease Activity via Long Range Resonance Energy Transfer, Journal of fluorescence, 23 (2013) 697–703. https://doi.org/10.1007/s10895-013-1189-7

[17] Q. Xu, Y. Zhang, C.-y. Zhang, A triple-color fluorescent probe for multiple nuclease assays, Chemical Communications, 51 (2015) 9121–9124. https://doi.org/10.1039/C5CC02177D

[18] I. Lehman, A. Nussbaum, The deoxyribonucleases of Escherichia coli V. On the specificity of exonuclease I (phosphodiesterase), Journal of Biological Chemistry, 239 (1964) 2628–2636.

[19] P.G. Mitsis, J.G. Kwagh, Characterization of the interaction of lambda exonuclease with the ends of DNA, Nucleic acids research, 27 (1999) 3057–3063. https://doi.org/10.1093/nar/27.15.3057

[20] Y. Zhang, Y. Wang, S.F.A. Rizvi, Y. Zhang, Y. Zhang, X. Liu, H. Zhang, Detection of DNA 3’-phosphatase activity based on exonuclease III-assisted cascade recycling amplification reaction, Talanta, 204 (2019) 499–506. https://doi.org/10.1016/j.talanta.2019.06.027

[21] C.C. Richardson, 16 Bacteriophage T4 Polynucleotide Kinase, The enzymes, Elsevier 1981, pp. 299–314. https://doi.org/10.1016/S1874-6047(08)60342-X

